# Metabolic reprogramming and altered ATP content impair neuroprotective functions of microglia in β-glucocerebrosidase deficiency models

**DOI:** 10.1101/2025.05.26.656111

**Authors:** Electra Brunialti, Alessandro Villa, Eva M. Szego, Pietro La Vitola, Denise Drago, Radmila Pavlovic, Laura Fontana, Doga Tuna, Alessia Panzeri, Clara Meda, Ornella Rondinone, Mattia Pitasi, Monica Miozzo, Annapaola Andolfo, Donato A. Di Monte, Paolo Ciana

## Abstract

Mutations in the *GBA* gene, which reduce β-glucocerebrosidase (GCase) activity, represent the most significant genetic risk factor for Parkinson’s disease (PD). Decreased GCase activity has also been observed in sporadic PD cases, supporting a broader role for GCase in the poorly understood mechanisms underlying PD etiopathogenesis. While most studies on the relationship between *GBA* mutations and PD have focused on neurons, evidence suggests that PD pathology promoted by GCase deficiency involves other cell types and, in particular, interactions between neuronal and glial cells.

Here, we identify microglia as primary players undergoing significant alterations at early stages of the pathological processes triggered by a GCase impairment. Using both pharmacological and genetic mouse models of GCase deficiency, we observed microglial morphological, transcriptional and metabolic changes. Interestingly, these changes were associated with a cell-specific, significant reduction of microglial ATP levels. When microglial ATP depletion was reproduced in an *in vitro* system of co-cultured microglial and neuronal cells, the neuroprotective properties of microglia were compromised and neuronal susceptibility to oxidative stress was enhanced.

These findings underscore the role of microglia in PD pathogenesis and point to a pathogenetic mechanism by which microglial metabolic disturbances leading to ATP depletion enhance neuronal vulnerability to injury and neurodegeneration. This mechanism could be targeted for therapeutic intervention aimed at mitigating PD risk and counteracting the development of PD pathology.

**GRAPHICAL ABSTRACT:** 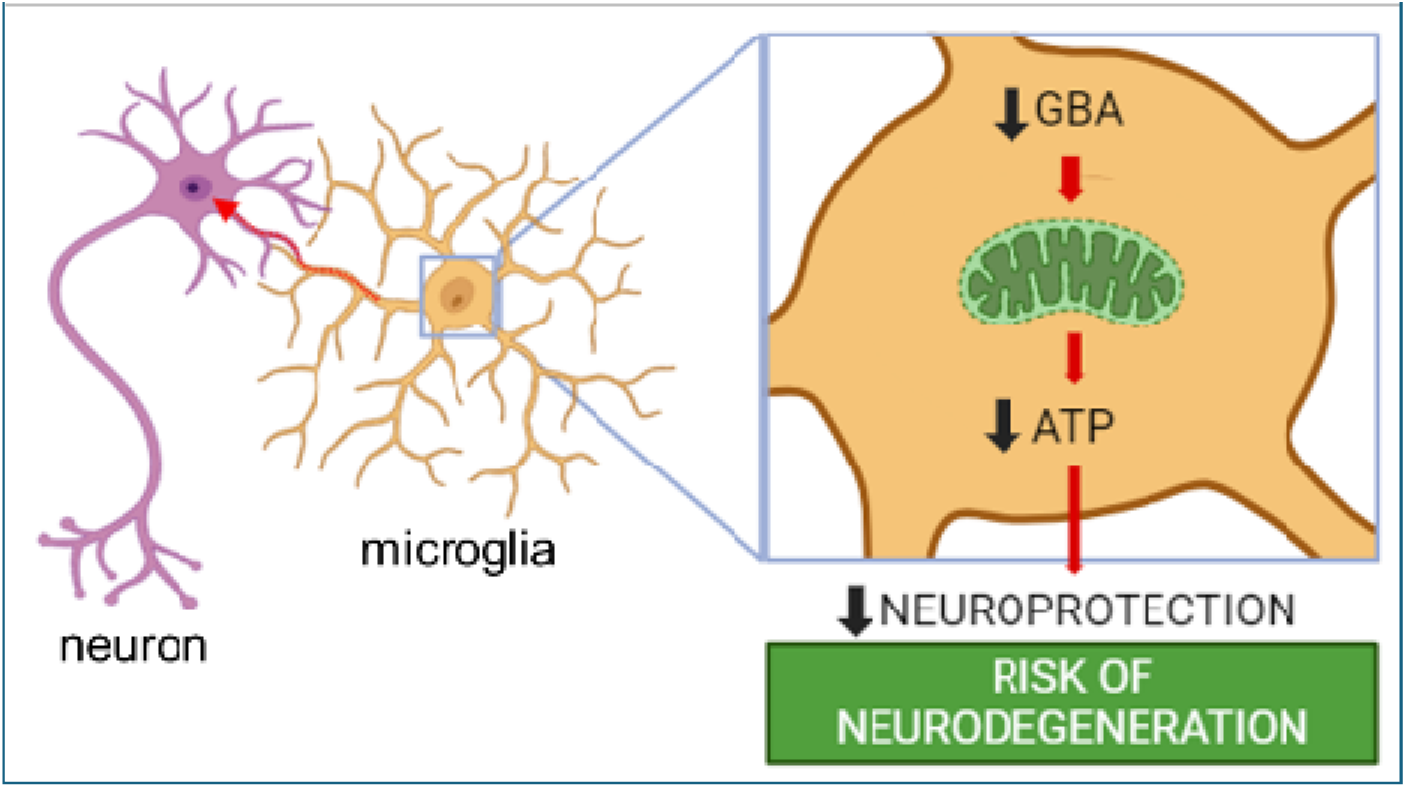

## Introduction

The human *GBA* gene encodes β-glucocerebrosidase (GCase), a lysosomal enzyme crucial for cellular function^1^. Mutations in both alleles of *GBA* significantly reduce GCase activity, leading to Gaucher’s disease (GD), a condition that primarily affects macrophages^2^. Notably, both homozygous and heterozygous *GBA* mutations are strongly associated with Parkinson’s disease (PD), with carriers facing a 20- to 30-fold increased risk^3^. These mutations can induce Parkinsonian symptoms that closely resemble idiopathic PD, indicating potential overlapping pathological pathways^4^. Interestingly, even PD patients without *GBA* mutations exhibit reduced GCase activity, suggesting that GCase dysfunction may play a broader role in PD pathogenesis^4,5^.

Despite extensive research, the mechanisms linking GCase impairment to neurodegeneration in PD remain not fully understood^4^. While most studies have focused on the impact of *GBA* mutations in neurons, evidence indicates that GCase deficiency in dopaminergic neurons alone does not fully explain the neuronal death and motor deficits seen in PD models^6^ . This suggests that pathological processes triggered by *GBA* mutations also involve other cell types. Several RNA sequencing studies in mice have shown higher *Gba* transcript levels in glia compared to neurons (brainrnaseq.org; holt-sc.glialab.org; astrocyternaseq.org). In this context, microglia, the brain’s resident immune cells, are particularly intriguing. Microglia maintain brain homeostasis through various functions, including immune surveillance, synaptic pruning, cellular debris clearance, and neurotransmission^7,8^. These activities demand substantial energy, requiring metabolic adaptation of microglia in response to different stimuli and conditions in order to maintain brain function and integrity; this adaptation involves, for example, switches between glycolysis and oxidative phosphorylation^9,10^.

In advanced stages of both Parkinson’s and Gaucher’s diseases, microglial activation and neuroinflammation are consistently observed, supporting a significant contribution of inflammation to neurodegenerative processes^11–16^. Notably, enhancing GCase activity using isofagomine reduces microglial inflammation and improves motor function in transgenic mice overexpressing α-synuclein^17^. Although microglia are known to play a pro-inflammatory role in later stages of the disease, emerging evidence suggests they also contribute to early pathological stages. As supports, in a zebrafish model of Gaucher’s disease, microglial activation occurs before neuronal death, implicating their involvement in the early stages of neurodegeneration^18^. Furthermore, positron emission tomography (PET) scans have identified microglial activation in regions vulnerable to Lewy body pathology in asymptomatic *GBA* mutation carriers^19^.

In line with this evidence, our previous research has shown that GCase inhibition in microglia leads to altered morphology and functionality, impairing their communication with neurons. These alterations hinder the activation of NFE2L2, a transcription factor that regulates redox balance and mitochondrial function by controlling the expression of antioxidant enzymes^20^. This disruption enhances neuronal susceptibility to a variety of toxic conditions/insults, supporting the hypothesis that an impairment of microglia-associated neuroprotective pathways may increase the risk of neuronal damage in PD and other human neurodegenerative diseases^21–23^ .

In this study, we aim to better characterize the neuroprotective phenotype of microglia and their role in the early pathogenetic stages by further investigating their phenotype in *GBA*-deficient models and elucidating the molecular mechanisms underlying impaired NFE2L2 mediated neuroprotection.

## Results

### Distinctive microglial morphology is present in GBA impaired models

The rationale for investigating changes in microglial morphology induced by a GCase impairment in the mouse brain was twofold. First, microglial morphology provides significant insight into microglial function and pathophysiology^24^. Second, when neuron-microglia interactions involving NFE2L2 were previously assessed *in vitro*, abnormalities caused by a GCase deficiency were found to be associated to overt microglial morphological changes. Here, we performed immunostaining of microglia in the brains of wild-type C57BL/6 mice following acute GCase inhibition. Six mice were divided into two experimental groups and treated with a daily intraperitoneal (i.p.) dose of 100 mg/kg/day of conduritol B epoxide (CBE), a GCase inhibitor, or vehicle (PBS) for three days. This treatment has been proven to effectively reduce microglial GCase activity, and to impact NFE2L2-mediated communication between microglia and neurons^21^.

Postmortem analyses were performed on tissue sections of the ventral mesencephalon, specifically targeting the substantia nigra pars compacta (SNpc), which were immunostained with anti-Iba1 antibodies (Figure 1A). The analysis of the stained microglia revealed a significant increase in microglia numbers in CBE-treated mice compared to vehicle-treated controls. Additionally, morphological examination identified notable alterations in the microglia of the CBE-treated group, which displayed a more ramified phenotype, as evidenced by an increased number of branches and triple points (junctions with exactly 3 branches) (Figure 1A, figure S1). Furthermore, these microglia exhibited a larger perimeter and greater extension, as indicated by the bounding box area increase (Figure 1A). However, no substantial differences were observed in branch length or solidity. Collectively, these observations indicate that microglial alterations are detectable at early stages following GCase inhibition.

**Figure 1:**
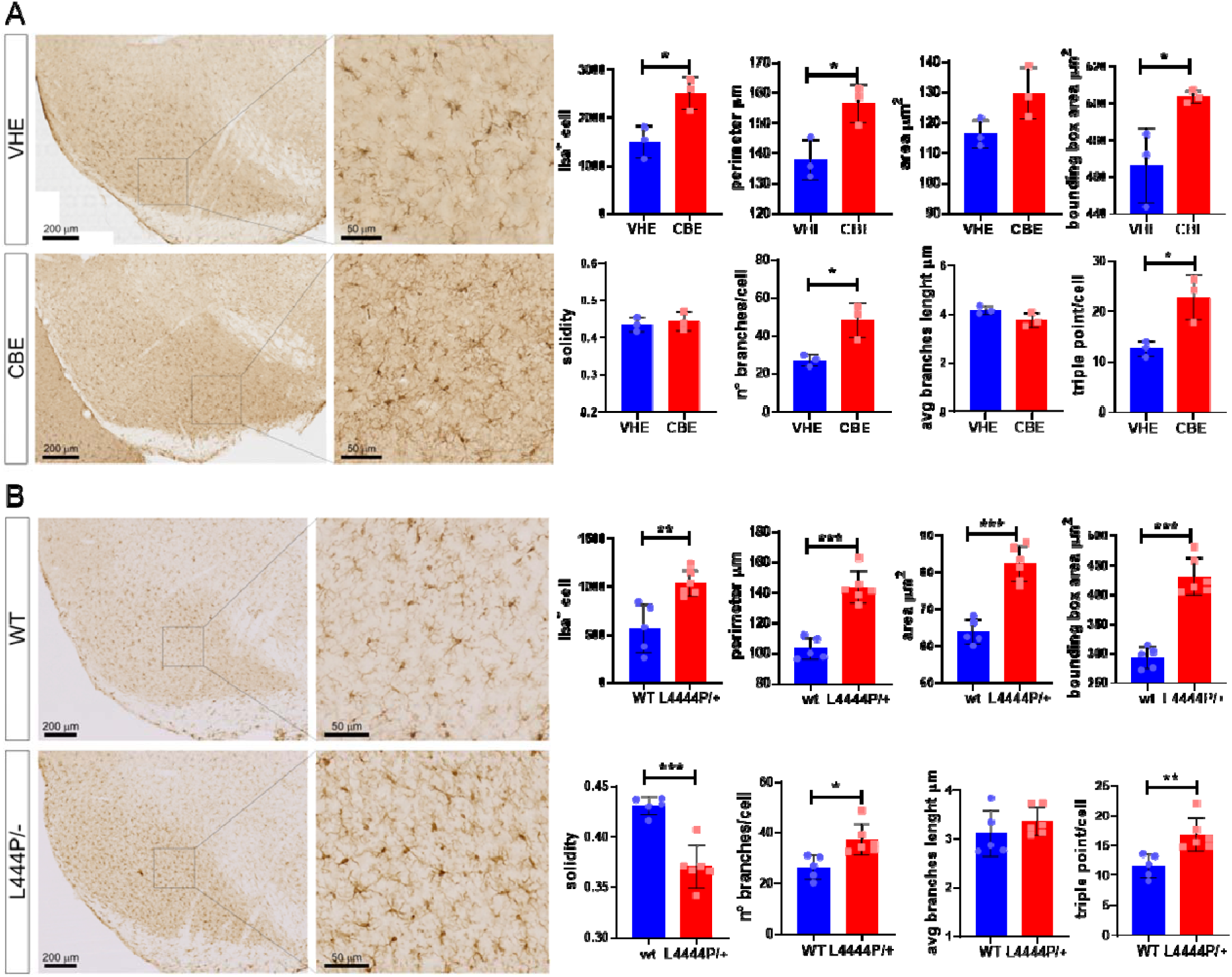
morphological characterization of GCase impaired microglia. (A) Representative image of the SNpc showing Iba1-positive microglia stained with DAB (left), along with an enlarged view (Scale bar, 200 µm and 50 µm) from vehicle-treated (VHE) and CBE-treated (CBE) mice (100 µg/kg for 3 days). Right, quantitative analysis of microglial morphology. Data are presented as average values per mouse expressed as mean ± SD. *p < 0.05 versus vehicle, determined using an unpaired t-test. *n* = 3 independent biological samples. (B) Representative image of the SNpc showing Iba1-positive microglia stained with DAB (left), along with an enlarged view (Scale bar, 200 µm) from L444P/+ and wt littermate along with quantitative analysis of microglial morphology (right). Data are presented as average values per mouse expressed as mean ± SD. *p < 0.05, **p < 0.01, ***p<0.001 versus wt, determined using an unpaired t-test. *n* = 5 for wt and *n*= 6 for L444P/+ independent biological samples.

Next, we tested whether altered microglial phenotypes were also present in a mouse genetic model bearing a PD-associated mutation in the *Gba* gene. We used B6;129S4-Gbatm1Rlp/Mmnc mice (hereafter referred to as L444P/+ mice) and their related wild-type control littermates. L444P/+ mice express a heterozygous knock-in L444P mutation in the murine Gba1 gene^25^. These mice exhibit a cerebral reduction of GCase activity of around 30-40%^26^. While aged mice do not display overt Parkinson’s disease-like behavioral changes, they show increased susceptibility to PD-relevant experimental conditions such as overexpression of human alpha-synuclein^27^. Therefore, they represent a suitable model for studying early pathophysiological changes that may enhance vulnerability to neurodegenerative processes.

Postmortem analyses were performed on tissue sections of the ventral mesencephalon, specifically the substantia nigra pars compacta (SNpc), from L444P/+ mice and control littermates (aged 55-60 weeks) (Figure 1B). Morphological analysis of Iba1-immunoreactive microglia in the SNpc revealed that the transgenic model exhibited an increased number of microglia with distinctive morphological features. These microglia were more highly ramified, larger, and less simple in structure, as indicated by an increase in branch number, area, bounding box area, triple points, and perimeter, alongside a decrease in solidity. These findings reveal significant microglial morphological alterations associated with heterozygous expression of L444P *Gba* in the mouse brain. Together with the results in CBE-treated animals, they also support an overall relationship between GCase deficiency and morphological microglial abnormalities.

### Identification of crucial pathways altered in GBA impaired models via transcriptomic analysis

To interrogate the cellular mechanisms associated to altered microglial morphology we conducted a transcriptomic analysis on microglia with inhibited GCase. Using CD11b microglia magnetic beads, primary microglia were isolated and purified from the brains of wild-type mice who received a daily intraperitoneal (i.p.) dose of 100 mg/kg/day CBE or vehicle (PBS) for 3 days. The transcriptome was investigated by RNA sequencing (RNA-seq). RNA was extracted from microglia pools collected from four brains each, with two pools derived from vehicle-treated mice and two pools from CBE-treated mice. Thus, a total of height mice per treatment were utilized in our study. Salmon was employed to analyze the transcriptional data count and obtain Transcripts Per Kilobase Million (TPM). Biomarkers representing neutrophils (Ly6g), B cells (Cd19), T cells (Cd3d, Cd8a, Cd8b1), astrocytes (Aldh1l1, Gfap), neurons (Thy1, Rbfox3), and oligodendrocytes (Olig1, Olig2) were found to be negligible in comparison to the predominant microglial markers (C1qa, P2ry12, Csfr1)(Figure S2) thus ascertaining microglia purity. Upon comparing CBE-treated microglia with controls, we identified 440 differentially expressed genes (DEGs) (Figure 2A) (P.Value < 0.05). To validate the reliability of the RNA-seq results, microglia were isolated from additional 16 mice. qPCR was performed on a panel of 9 mRNAs selected from those exhibiting treatment-dependent or independent expression, confirming the reproducibility of the RNA-seq data (Figure S3).

**Figure 2:**
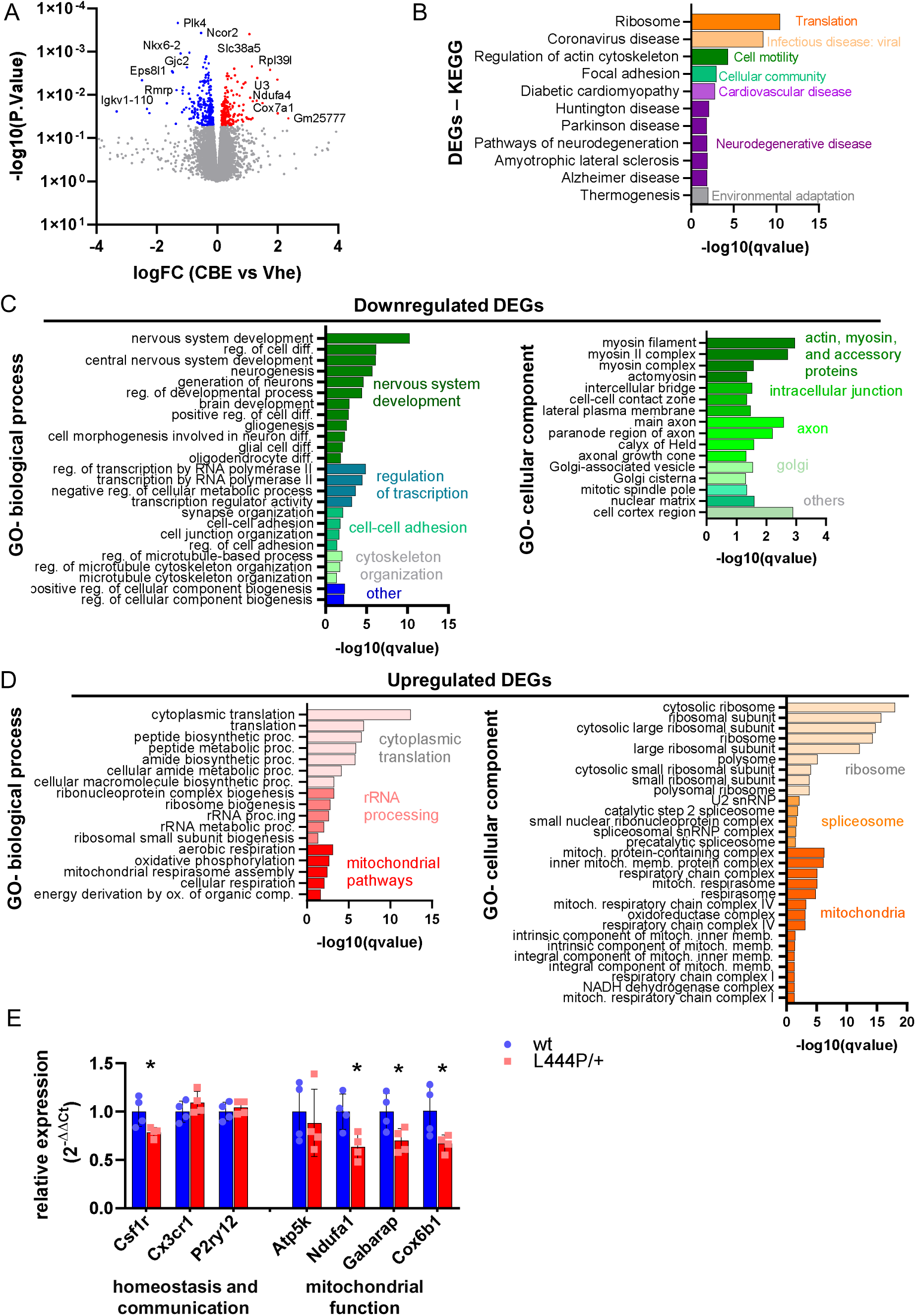
Microglia with impaired GBA function exhibit a distinct transcriptomic signature. (A) Volcano plot of RNA-seq data obtained from microglia isolated from mice treated with 100 mg/kg/day CBE for three days. Analyses were conducted on two pools of four brains each (total of 8 mice per condition). Blue dots represent genes with significantly lower TPM values in CBE-treated microglia compared to vehicle-treated microglia. Red dots represent genes with significantly higher TPM values in CBE-treated microglia compared to vehicle-treated microglia (p < 0.05). An unpaired t-test was used as the statistical test. (B) Pathways analysis of differentially expressed genes (DEGs) using the Kyoto Encyclopedia of Genes and Genomes (KEGG) databases. Significance, calculated with two-sided hypergeometric test and corrected with the Bonferroni step-down method, is represented on the x axis (–log10 scale of the q-value). Pathways are grouped and colored according to the BRITE hierarchy class. (C) Gene Ontology analysis of DEGs referring to biological processes (left) and cellular components (right) of (C) downregulated and (D) upregulated DEGs. Significance, calculated with two-sided hypergeometric test and corrected with Bonferroni step-down, is represented on the x axis (–log10 of the q-value). Pathways are grouped and colored by biologically related processes. (E) Modulation of gene expression in microglia purified from L444P/+ mice. Total RNA was purified from L444P/- mice, and the expression of selected mRNA genes related to homeostasis, microglia-neuron communication, and mitochondrial functionality was analyzed by real-time PCR. Relative quantification of the transcripts was obtained using the 2−ΔΔCt method versus wt littermates. Data are presented as mean ± SD; *p<0.05 determined by t-test. Analyses were conducted on four pools of two brains each.

To investigate whether the differentially expressed genes (DEGs) were associated with specific functional categories within large gene clusters, we performed a cluster analysis using Cytoscape with the ClueGO plug-in^28^. Statistical analysis was conducted using a two-sided hypergeometric enrichment/depletion test, with Bonferroni step-down correction applied for multiple testing. A comprehensive analysis of all DEGs (both upregulated and downregulated) through the Kyoto Encyclopedia of Genes and Genomes (KEGG) pathway analysis (Figure 2B) highlighted that the affected pathways were mainly associated to translation, cell motility and communication and various neurodegenerative diseases, including Parkinson’s disease. In our examination of downregulated genes (Figure 2C), the functional analysis of Gene Ontology (GO) biological processes revealed a predominant association with pathways related to neuronal development, regulation of transcription, cell adhesion, and cytoskeleton organization. Furthermore, the GO cellular components analysis showed an enrichment of proteins associated with the contact zone between cells, actin-myosin filaments, axons, and Golgi-associated vesicles. These findings align with the previously identified phenotype of microglia following GCase inhibition, wherein the cells exhibit impaired motility^23^ and morphology (Figure 1A) and affected communication to the neurons^21^.

Concerning the upregulated genes, the GO functional analysis indicated that the treatment led to an increase in pathways related to cytoplasmic translation, rRNA processing, and energy metabolism generated by mitochondria and cellular respiration (Figure 2D). The cellular component analysis (Figure 2D) further revealed that upregulated genes were predominantly associated with the ribosome, spliceosome, and mitochondrial cellular compartments. Notably, we observed an enrichment in proteins involved in the mitochondrial respiratory chain.

Next, we investigated whether L444P/+ microglia exhibit similar alterations in genes associated with homeostatic microglial functions and mitochondrial activity. This analysis of the relevant genes identified with the RNAseq experiment was performed using real-time PCR on mRNA purified from microglia isolated from L444P/+ mice, with comparisons made to their wild-type littermates (Figure 2E). mRNA samples from microglia of two brains were pooled, resulting in four pools per group for analysis. The results revealed a reduction in the expression of genes associated with both microglial homeostasis and mitochondrial functionality. Specifically, Csf1r, Ndufa1, Gabarap, and Cox6b exhibited statistically significant reductions in expression levels. These findings suggest that microglia transcript alterations are also present in this genetic model, further supporting the notion that they are a consequence of both pharmacologic and genetic GCase deficiency.

### GCase impairment affects microglial metabolism

Since transcriptomic analysis suggested that metabolic alterations are induced in both the studied models, we further investigated the cellular changes induced by GCase inhibition by conducting metabolic profiling of CBE-treated microglia. Mice treatment and microglia purification were performed as described previously. For this study, groups of six brains were pooled, and metabolites were extracted from four pools per condition. We quantified the metabolites associated with key energetic pathways, including glycolysis, the pentose phosphate pathway (PPP), the tricarboxylic acid (TCA) cycle, and glutaminolysis (Figure 3A).

**Figure 3:**
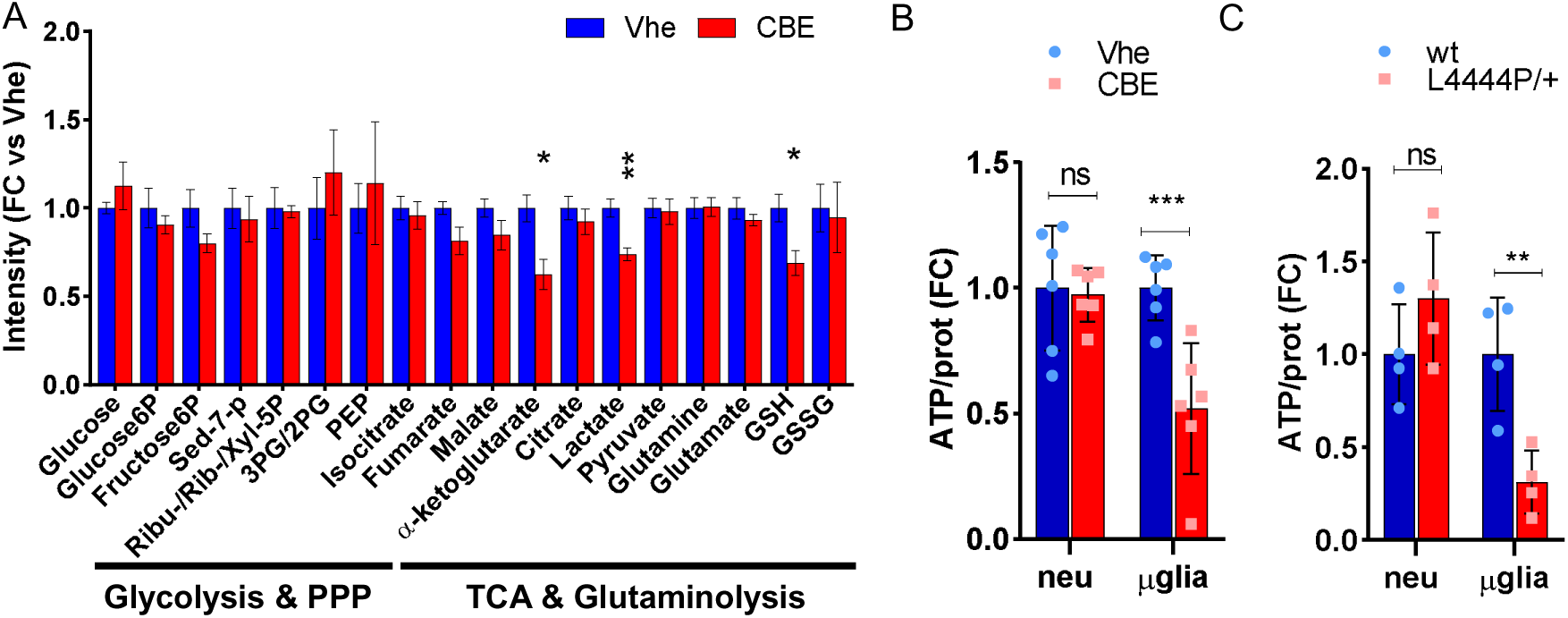
Metabolic and energetic alterations in GBA impaired microglia. (A) Metabolic profile of microglia purified from mice treated with 100 mg/kg/day CBE for three days. Analyses were conducted on four pools of six brains each (total of 24 mice per condition). Data represent fold change (FC) of metabolite intensity compared to vehicle ± SEM. *p < 0.05, **p < 0.01 versus vehicle, calculated with unpaired t-test. PPP = Pentose Phosphate Pathway; TCA= Tricarboxylic Acid Cycle; Sed-7-p = Sedoheptulose-7-phosphate; Ribu-/Rib-/Xyl-5P= Ribulose-5P/Ribose-5P/Xylulose-5P; 3PG/2PG = 3-e 2-Phosphoglycerate; PEP = Phosphoenolpyruvate; GSH = reduced glutathione; GSSG = oxidized glutathione. (B) Relative quantity of ATP detected in microglia cells and in neuronal populations extracted from mice treated with 100 mg/kg/day CBE for three days. Data represent fold change (FC) of ATP amount normalized to total protein extract versus vehicle ± SD (n=6); ***p < 0.001 versus vehicle, calculated with one-way ANOVA followed by Sidak’s multiple comparisons test. Neu= mixed brain cells population lacking microglia; µglia= microglia. (C) Relative quantity of ATP detected in microglial cells and neuronal populations extracted from L444P/+ and wt littermates. Data represent FC of ATP amount normalized to total protein extract versus vehicle ± SD (n=4); **p<0.01 versus wt, calculated with one-way ANOVA followed by Sidak’s multiple comparisons test.

Among the identified metabolites, microglia with inhibited GCase showed significant reductions in lactate, reduced glutathione (GSH), and alpha-ketoglutarate, a TCA cycle intermediate that plays a critical role in immune cell function^29–31^.

Given the metabolomic and transcriptomic evidence suggesting mitochondrial and metabolic alterations, we also measured cellular ATP levels using a bioluminescent biochemical assay in microglia purified from CBE-treated mice (Figure 3B). Microglia with inhibited GCase showed a drastic reduction in ATP levels (-50%) compared to vehicle-treated controls, indicating that GCase deficiency reduces microglial ATP availability. To determine if this ATP alteration was specific to microglia or detectable in other brain cells, ATP content was also measured in a brain cell fraction enriched in astrocytes, neurons, and oligodendrocytes extracted from CBE-treated mice (Figure 3B, Figure S4). No alterations in ATP concentration were detected in the overall brain cell fraction between vehicle and CBE treatment, indicating that microglial mitochondrial activity is more impaired than that of other brain cells at early stages following acute CBE inhibition. Similar findings were observed in cells isolated from the brains of L444P/+ mice. ATP measurements in cellular extracts revealed a specific reduction in ATP levels in microglia from L444P/+ mice compared to their wild-type littermates, whereas no significant changes were observed in the mixed brain cell population lacking microglia (Figure 3C).

### Microglial ATP plays a key role in neuroprotective microglial communication

To further investigate the metabolic alterations observed and their correlation with microglial functionality, we shifted to a more standardizable and convenient *in vitro* system using immortalized cells. Our prior work demonstrated that co-cultures of immortalized dopaminergic neuroblastoma cells (SK-N-BE) and microglia (BV-2) provide an effective model for studying microglia-to-neuron communication, mirroring *in vivo* cross-talk^21,23^. We first measured cellular ATP content following GCase inhibition in immortalized BV-2 and SK-N-BE cells. Both cell types were treated with 200 µM CBE for 48 hours, a timeframe previously determined to almost completely abolish GCase activity and reduce microglia-to-neuron communication without affecting cell viability^21^. Cellular ATP levels were then quantified using a bioluminescent biochemical assay. Consistent with prior findings, only BV-2 microglia exhibited a significant reduction in ATP levels following CBE treatment (about -20%), while SK-N-BE cells showed no significant differences in ATP levels between vehicle and treated groups (Figure 4A).

**Figure 4:**
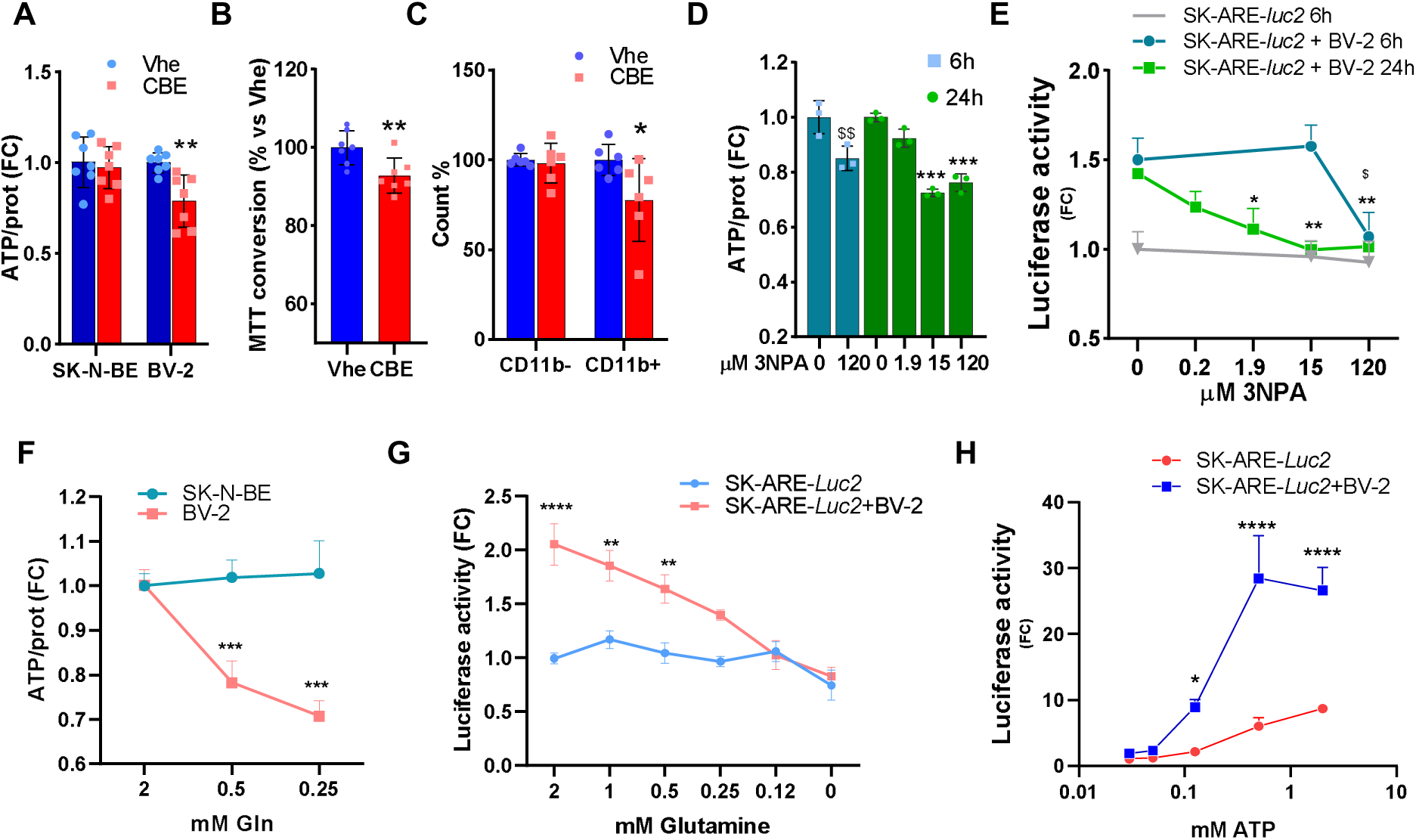
Microglial ATP plays a central role in microglia-to-neuron communication. (A) Relative quantity of ATP detected in SK-N-BE and BV-2 cells treated with vehicle or 200 µM CBE for 48 hours. Data represent fold change (FC) of ATP amount normalized to total protein extract versus vehicle ±SD (n=7); **p<0.01 versus vehicle, calculated with one-way ANOVA followed by Sidak’s multiple comparisons test. (B) Conversion of MTT in co-cultures of SK-N-BE and BV-2 cells treated with vehicle or 200 µM CBE for 48 hours. Data represents percentage of substrate conversion relative to vehicle-treated cells ±SD (n=7); **p< 0.01 versus vehicle, calculated with unpaired t-test. (C) Flow cytometry assay showing the mitochondrial intensity staining of CD11b-cells and CD11b+ cells co-cultured and treated with vehicle or 200 µM CBE for 48 hours. Data represents percentage of stained cells ±SD (n=7); *p< 0.05 versus vehicle, calculated with one-way ANOVA followed by Sidak’s multiple comparisons test. (D) Relative quantity of ATP detected in BV-2 cells treated with vehicle or different concentrations of 3-nitropropionic acid (3-NPA) for 6 or 24 hours. Data represent FC of ATP amount normalized to total protein extract compared to vehicle ±SD (n=3); ^$$^p<0.01 versus vehicle at 6 hours, ***p<0.0001 versus vehicle at 24 hours, calculated with one-way ANOVA followed by Dunnett’s multiple comparisons test. (E) SK-ARE-*luc2* cells or coculture of SK-ARE-*luc2* and BV-2 cells were treated with 200 μM 3-NPA for 6 or 24 hours, and bioluminescence representative of NFE2L2 activation in SK-ARE-*luc2* cells was measured as a marker of microglia-to-neuron communication. Data are expressed as fold FC of RLU versus vehicle and presented as mean ±SEM (n=8); *p<0.05, **p<0.01 versus vehicle for 24 hours treatment, ^$^p<0.05 versus vehicle for 6 hours treatment calculated with one-way ANOVA followed by Dunnett’s multiple comparisons test. (F) Relative quantity of ATP detected in SK-N-BE and BV-2 cells grown in complete media with different concentrations of glutamine for 6 hours. Data represent FC of ATP amount normalized to total protein extract compared to vehicle ±SEM (n=8); ***p<0.001 versus SK-N-BE calculated with one-way ANOVA followed by Sidak’s multiple comparisons test. (G) Luciferase activity was measured in protein extracts from SK-ARE-*luc2* cells or cocultures of SK-ARE-*luc2* and BV-2 cells, treated for 4 hours with increasing doses of ATP. Data represents the FC in luciferase activity compared to vehicle-treated SK-ARE-luc2 cells, presented as mean ±SEM (n=10). *p < 0.05, ****p<0.0001 versus SK-ARE-luc2, as determined by one-way ANOVA followed by Sidak’s multiple comparisons test. (H) Luciferase activity measured in protein extracts derived from SK-ARE-*luc2* cells or coculture of SK-ARE-*luc2* and BV-2 cells grown for 6 hours in complete media containing different concentrations of glutamine. Data represent FC of luciferase activity versus SK-ARE-*luc2* grown in complete media ±SEM (n=18); **p<0.01, ****p<0.0001 versus SK-ARE-*luc2* calculated with one-way ANOVA followed by Sidak’s multiple comparisons test.

To investigate whether this ATP reduction was linked to altered metabolic activity, we assessed central metabolic pathway functionality using the MTT colorimetric assay. BV-2 and SK-N-BE cells were co-cultured and treated with 200 µM CBE for 48 hours. Cell viability, confirmed via trypan blue exclusion (Figure S5), was not affected; however, formazan production was reduced by approximately 10% in CBE-treated co-cultures compared to controls (Figure 4B), indicating altered metabolic activity.

Next, to determine if microglial mitochondria were specifically affected by GCase inhibition, we stained co-cultured cells treated with CBE using MitoTracker Red, which accumulates in active mitochondria, and CD11b, a microglial marker. Flow cytometry analysis (Figure 4C, Figure S6) revealed reduced mitochondrial fluorescence specifically in the CD11b+ population (BV-2 cells), suggesting specific mitochondrial impairment in microglia due to GCase inhibition.

Based on our previous findings showing that GCase inhibition impairs neuroprotective microglia-to-neuron communication^21^, we hypothesized that mitochondrial dysfunction of microglia underlies this impaired neuroprotection. To test this hypothesis, we used a co-culture system where neuroblastoma (SK-N-BE) cells stably transfected with a reporter vector of NFE2L2 activity (named SK-ARE-*luc2*) were cultured with BV-2 microglia^21^. This model effectively assesses, in a semi-quantitative way, the strength of microglia-to-neuron communication by measuring bioluminescence from the neuronal reporter cells (Figure S7A)^21^.

To induce mitochondrial dysfunction and reduce ATP content, we treated BV-2 cells with 3-nitropropionic acid (3-NPA), an inhibitor of succinate dehydrogenase that impacts both the TCA cycle and the mitochondrial respiratory chain. Treatment with 3-NPA at 120 µM for 6 hours or 15– 120 µM for 24 hours resulted in a 20–25% reduction in ATP levels in BV-2 cells compared to controls (Figure 4D). In co-cultures (SK-ARE-*luc2* + BV-2) treated with 3-NPA, a dose-dependent reduction in neuronal NFE2L2 activation was observed, correlating with ATP depletion (Figure 4E) supporting the potential link between energy metabolism and microglial neuroprotective functions. Similar results were obtained by reducing ATP via starvation growing the cell for six hours in a nutrient-depleted media (75%, 50%, and 25% complete medium) (Figure S7B).

It is known that homeostatic microglia predominantly rely on oxidative phosphorylation (OXPHOS) and glutaminolysis for ATP production, whereas neurons primarily depend on glucose metabolism^9,32–34^. Therefore, to investigate whether reducing glutamine levels selectively impacts microglial ATP content, we cultured BV-2 and SK-N-BE cells in media containing reduced glutamine concentrations (0.5–0.25 mM) for 6 hours. This treatment significantly decreased ATP levels in BV-2 cells, with no notable effects on SK-N-BE cells (Figure 4F), indicating that lowering glutamine is an effective approach to selectively reduce ATP in microglia. Next, to test the role of microglial ATP in their communication with neuron, co-cultures of SK-N-BE reporter and BV-2, as well as SK-N-BE reporter cells alone, were subjected to the same treatment. Despite no detectable loss in cell viability (Figure S7C), co-cultures exhibited reduced neuronal NFE2L2 activation, coinciding with diminished microglial ATP levels (Figure 4F, 4G). These findings suggest a direct role of microglial ATP in their neuroprotective communication with neurons. Similarly, treatment with the glutamine analog 6-diazo-5-oxo-L-norleucine (DON), an inhibitor of glutamine-utilizing enzymes, produced comparable effects (Figure S7D).

Finally, we tested whether ATP supplementation could restore microglia-to-neuron communication. A method to selectively activate OXPHOS and induce intracellular ATP accumulation in microglial cells is to activate purinoreceptors via ATP addition^35,36^. Indeed, the addition of ATP to the neuron-microglia co-culture elicited a rapid increase in microglia-to-neuron communication, evidenced by a clear dose-dependent increase in the reporter signal detected 4 hours after ATP addition (Figure 4H). Conversely, the addition of pro-inflammatory stimuli known to promote glycolysis while suppressing OXPHOS and inducing mitochondrial fragmentation, such as lipopolysaccharide (LPS)^37,38^, did not affect microglia-to-neuron communication (Figure S7E). This finding corroborates the role of ATP and mitochondrial functionality in microglia-to-neuron communication.

## DISCUSSION

Decreased GCase activity has been observed in GBA carriers as well as in the brains of patients with idiopathic PD^39^, a finding that highlights the significance of studying GCase dysfunction to gain deeper insights into the etiopathogenesis of PD. Research on the role of GBA mutations in PD has predominantly focused on the direct effects of altered GCase in dopaminergic neurons; however, accumulating evidence suggests that glial GCase deficiency, particularly in microglia, contributes significantly to disease pathogenesis^21,40–42^. Although immune dysregulation is well documented in Gaucher disease ^43^, the involvement of microglia during the early phases of GBA-associated PD remains to be fully clarified. Notably, microglial activation has been detected in Lewy body– vulnerable regions of asymptomatic GBA mutation carriers^19^ and in the brains of GCase-deficient zebrafish models of Gaucher disease^18^, suggesting that microglial dysfunction may precede manifest neuronal loss. Our previous work demonstrated that impairing microglial GCase activity reduces their neuroprotective capabilities, thereby increasing neuronal susceptibility to PD-related stressors^21,23^. Collectively, these observations imply that dysfunctional microglia could play a role as a mechanism involved in early stages of PD pathological processes.

In the present study, we extended these findings by characterizing the microglial phenotype induced by GCase impairment. Both the chemically induced (via CBE treatment) and genetic (L444P/+ mutant *Gba*) models of GCase loss exhibited early microglial alterations, including increased cell numbers and enhanced ramification, indicative of an activated state. These morphological changes occurred in the absence of any overt evidence of behavioral or neurodegenerative abnormalities, supporting the conclusion that microglial activation was not a mere consequence of tissue damage but actually preceded and may possibly play a “causative” role in pathological processes associated with GCase deficiency. Transcriptomic analysis of microglia identified that genes involved in cytoskeleton organization, cell–cell adhesion, and neuroprotective signaling pathways. Alterations in these pathways likely play a role in both morphological and functional changes that affect microglial cells with lowered GCase activity. As importantly, changes in these pathways could significantly affect microglia-neuron interactions and impair the critical function of microglia capable of monitoring, protecting and rescuing neurons through specialized junctions and structural nanotube connections^43,44,45^ (Figure 1). Altered cytoskeleton organization and neuroprotective signaling pathways would also underlie the results of earlier investigations that specifically showed paralleled changes in microglial morphology and microglia-related neuroprotection as a consequence of GCase inhibition^21^.

Our transcriptomic data also revealed changes in the expression of genes associated with protein synthesis, RNA processing, and mitochondrial function. Interestingly, similar gene expression alterations have been identified in peripheral blood mononuclear cells from PD patients irrespective of GBA mutation status^46^, suggesting myeloid cell dysregulation as a hallmark of PD, which, in the microglial lineage, contributes to neuronal susceptibility to pathogenetic insults. The conservation of these alterations across various forms of PD highlights their potential relevance in disease pathogenesis.

Metabolic profiling further revealed that GCase inhibition in microglia leads to distinct perturbation in microglial metabolism. In particular, we observed significant reductions in metabolites such as lactate, reduced glutathione (GSH), and α-ketoglutarate. Since the shift between glycolysis and oxidative metabolism produces phenotypic and functional changes in microglia^47^, this suggests that investigating the link between mitochondrial function, α-ketoglutarate, glutamine, and energy metabolism could provide valuable insights into microglial energy balance and impairment of its function during disease. In detail: (i) the reduced levels of lactate that we observed after GCase inhibition in microglia may indicate an impairment of the glycolytic flux^48^; (ii) the decrease in GSH is indicative of increased oxidative stress and impaired cysteine metabolism due to the accumulation of glucosylceramide and related lipids^49^; and (iii) the reduced α-ketoglutarate levels may result from disruptions in the TCA cycle, as GCase inhibition can impair mitochondrial function, leading to decreased metabolic activity and α-ketoglutarate production^50^.

Furthermore, α-ketoglutarate plays a crucial role in glutaminogenesis, and its reduction may impair glutamine synthesis, which is essential for maintaining adequate ATP levels in microglia (Figure 4). This is a key finding of the present work: the ATP-related metabolic alterations resulting from GCase impairment are observed specifically in the microglial lineage, suggesting a cell-type-specific vulnerability. This phenomenon could be related to their high phagocytic activity and reliance on lysosomal lipid metabolism, resulting in glucocerebroside accumulation, lysosomal dysfunction, and secondary mitochondrial impairment.

Mitochondrial dysfunction is a well-established hallmark of both sporadic and GBA-associated PD. Our study confirms that GCase impairment leads to reduced mitochondrial staining and significant ATP depletion in microglia from both CBE-treated and L444P/+ models. Considering that microglia rely heavily on ATP for functions such as motility, surveillance, phagocytosis, and cytokine production, this energy deficit likely undermines their neuroprotective capacity. Experiments using succinate dehydrogenase inhibitors to mimic mitochondrial dysfunction further demonstrated that ATP depletion correlates with impaired microglia-to-neuron communication (Figure 4). In contrast, restoration of ATP levels through purinergic receptor activation improved this communication, underscoring the critical role of energy metabolism in sustaining microglial function^35,36^. These changes compromise microglial ATP production and, by extension, their ability to support neuronal survival, thereby potentially contributing to the increased neurodegenerative risk observed in PD-GBA.

Taken together, these findings strongly point to microglial changes that precede and are likely to play an important role in the sequence of toxic/pathological events underlying PD development. Our hypothesis is that the observed reduction of ATP production in microglia could represent a converging point of multiple pathogenic factors, including genetic predispositions and metabolic toxins known to affect mitochondrial functionality and, in turn, trigger PD-relevant neurodegenerative processes^51^. Despite diverse origins, these factors ultimately converge on mitochondrial dysfunction in microglia, leading to impaired ATP production. A deficiency in microglial ATP may, in turn, compromise neuroprotection, exacerbating neuronal vulnerability to injury and contributing to neuronal demise. In particular, loss of microglial ATP may impact on the important role that microglia-mediated NFE2L2 activation has in modulating neuronal oxidative stress response.

Mitochondrial impairment has long been suggested to play an important role in PD pathogenesis, and this hypothesis has found significant support from evidence in PD brain^27^. Interestingly, earlier work has attributed the pathological consequences of mitochondrial abnormalities in PD primarily to metabolic/toxic changes affecting neuronal cells. Our present findings provide a broader perspective, underscoring the pathogenetic relevance of changes in mitochondrial energy metabolism affecting microglia. Although microglial mitochondrial impairment was mostly assessed here in relation to GCase deficiencies, the previous considerations underscore the likelihood that microglial mitochondrial dysfunction and reduced ATP availability in microglia may not only be relevant for *GBA*-associated but also other forms of PD.

From a pharmacological point of view, targeting microglial energy metabolism might represent an effective strategy to mitigate PD risk and to counteract PD neurodegenerative processes. The ATP deficiency observed in GCase-impaired microglia suggests that restoring mitochondrial function could also restore their neuroprotective capacity. For instance, agents that enhance oxidative phosphorylation, such as coenzyme Q10, nicotinamide riboside, MitoQ, and elamipretide ^52–54^, or drugs that activate purinergic receptors to boost ATP synthesis, such as 2-methylthioadenosine 5′-triphosphate and benzoylbenzoyl-ATP^55,56,57^, may normalize microglial activity and improve neuron–microglia communication. In parallel, the altered metabolic pathways identified, including reductions in α-ketoglutarate and glutathione, offer additional targets for intervention, potentially attenuating the inflammatory environment that exacerbates neuronal vulnerability. The shift of cellular targets from neurons to microglia opens new therapeutic opportunities and warrants further evaluation as it may lead to alternative strategies for the treatment not only of patients with *GBA* mutations but also patients with idiopathic PD.

## Conclusion

The results presented in this study challenge the traditional paradigm that focuses on neurons as targets of metabolic/toxic alterations in PD and as key players in the development of parkinsonian pathology in carriers of *GBA* mutations. This study not only supports the emerging view ^21,40–42^ of a crucial role of microglia in the pathogenesis of neurodegenerative diseases, but also identifies a specific new mechanism of microglia-associated neuroprotection. In particular, data reveal that microglial mitochondrial impairment, which can arise from a variety of genetic, toxic or metabolic conditions, can negatively affect the ability of microglia to promote and sustain neuronal stress responses, thus enhancing neuronal vulnerability to injury. Loss of microglial ATP plays a key role in mediating this deleterious effect of abnormal neuron-microglia interaction. As a corollary to these findings, we propose that targeting altered microglial metabolism may represent an effective therapeutic strategy. By reverting changes in microglial metabolism, we could ultimately enhance neuronal resilience, thus reducing PD risk and counteracting disease development.

## Material And Methods

### Cell cultures

All cell lines were purchased from the American Type Culture Collection (ATCC). SK-ARE-*luc2* cells were obtained by stable transfection of SK-N-BE cells with pARE-luc2-ires-tdTomato plasmid^22^. For coculture experiments 70,000 SK-N-BE or SK-ARE-*luc2* cells were plated in each well of a 24-well plate and cultured for 1 day. Then 3,500 BV-2 cells/well were seeded for over the neuron layer. Cells were grown in Neurobasal A medium (Cat. 10888-022, Life Technologies) containing 1% streptomycin–penicillin, 1% GlutaMAX (Car. 35050061, Life Technologies), 2% B-27 Supplement (Cat. 17504-044; Gibco), and 10 mM HEPES (Cat. H0887, Merck) in a humidified 5% CO2/95% air atmosphere at 37 °C.

### Cell treatments

Cell treatments were performed as follows, unless otherwise specified. Cells were treated with 200 µM conduritol-B-epoxide (CBE, Cat. 234599, Merck) or vehicle (water) for 48 hours. For experiments involving 3-Nitropropionic acid (3-NPA, Cat. N5636, Merck), cells were treated for either 6 hours or 24 hours, with vehicle (water) as a control. ATP (Cat. A7699, Merck) was administered at a concentration of 200 µM for 4 hours, with water serving as the vehicle control. Cells were treated with 200 ng/ml LPS O111:B4 (Cat. L2630, Merck) for 6 hours, with vehicle (water) as the control. For the nutrient-depletion experiments, complete media was prepared with varying concentrations of GlutaMAX, which included final dosages of 2 mM, 1 mM, 0.5 mM, 0.25 mM, 0.12 mM, or no glutamine, as indicated in the specific experimental conditions.

### Flow cytometry assay

Flow cytometry experiments were performed on at least 200,000 cells for each sample by using a Novocyte 3000 (Agilent Technologies, Inc.) equipped with 405 and 640 nm lasers. Cells were incubated for 10 min with 45 ng of CD11b-VioBlue antibody (Cat. 130-113-810, Mitlenyi biotech) and 1:4000 MitoTracker Deep Red FM (Cat. M46753, Invitrogen). Fluorescence pulses were detected using a 445/45 and 675/30 nm collection filter. The results were analyzed using NovoExpress software (Agilent Technologies).

### Microglia and brain cells isolation

Microglia were isolated from the whole brains of adult (age 3-6 months) male mice using a protocol previously described by Brunialti et al. ^21^. In brief, the brains were subjected to both enzymatic and mechanical dissociation. Following myelin removal, the cell suspension was processed for sorting using a magnetic column (Cat. 130-093-634, Miltenyi Biotec) to isolate CD11b+ cells, which were identified as microglia. The suspension of cells not retained in the column, which lacked CD11b expression (CD11b-), was used as a neuronal population devoid of microglia.

### Luciferase enzymatic assay

Luciferase assays were performed as described previously ^21^. Briefly, cells were lysed with Luciferase Cell Culture Lysis Reagent (Cat. E1531, Promega), and the protein concentration was determined with a Bradford assay. The biochemical luciferase activity assay was carried out in luciferase assay buffer by measuring luminescence emission with a luminometer (Veritas, Turner Biosystems), and the relative luminescence units (RLU) were determined during 10-second measurements.

### ATP Detection

ATP levels were measured using the Luminescent ATP Detection Assay Kit (Cat. ab113849, Abcam) according to the manufacturer’s instructions. Briefly, cells were lysed using the provided detergent, followed by the addition of the Substrate Solution. Bioluminescence was measured using a luminometer (Veritas, Turner Biosystems), and ATP concentrations were quantified using a standard ATP calibration curve. Protein concentrations were determined using the Bradford assay.

### Animal

All animal experiments were carried out in accordance with the guidelines of the Federation for European Laboratory Animal Science Associations (FELASA) and approved by the State Agency for Nature, Environment and Consumer Protection in North Rhine Westphalia (Landesamt für Natur, Umwelt und Verbraucherschutz Nordrhein-Westfalen), Germany, project 84-02.04.2017.A211 and by the Italian Ministry of Research and University (permission numbers: 5247B.N.459/2019, 12-12-30012012/2012). The animals were housed in individually ventilated plastic cages, maintained within a temperature range of 22–25 °C and a relative humidity of 50% ± 10%. They were kept on a 12-hour light/dark cycle, with lights on at 07:00. Animals had free access to pelleted food and fresh tap water, and all cages were equipped with nesting material. For pharmacological treatments, wild-type C57BL/6 male mice (15–30 weeks old) were administered 100 mg/kg/day of conduritol-B-epoxide (CBE) or vehicle (PBS) via intraperitoneal (i.p.) injection for 3 consecutive days. For the genetic GBA model, heterozygous knock-in L444P mice (B6;129S4-Gbatm1Rlp/Mmnc) and non-transgenic littermates were obtained from the Mutant Mouse Regional Resource Centre (RRID: MMRRC_000117-UNC). L444P/wt mice were identified by PCR of the genomic DNA using the following forward and reverse primers: 5’-CCCCAGATGACTGATGCTGGA-3’ and 5’-CCAGGTCAGGATCACTGATGG-3’ (dx.doi.org/10.17504/protocols.io.5qpvo3xobv4o/v1). PCR amplification products were digested by the restriction enzyme *NciI*, yielding two fragments at 386 and 200 bp in heterozygous animals. 40–50 weeks old male mice were used in the assays.

### Microglia staining in brain sections and analysis

Mice were sacrificed via intraperitoneal (i.p.) injection of sodium pentobarbital and transcardially perfused with cold saline solution followed by 4% paraformaldehyde in PBS. The brains were immediately removed, postfixed in 4% paraformaldehyde for 24 hours and then cryopreserved in 30% sucrose. Serial coronal sections of 35 μm were cut throughout the brain using a freezing microtome. To stain microglia in the substantia nigra, immunohistochemistry with brightfield detection was performed in serial coronal brain slices encompassing one-tenth of the whole substantia nigra. Free-floating sections were incubated in a quenching solution (3% H2O2 and 10% methanol in TRIS-buffered saline, pH 7.6). Non-specific binding was blocked by incubating samples in 5% normal goat serum. Iba1 was detected using a specific rabbit anti-Iba1 primary antibody (1:5000; Wako, Cat. 019-19741). After incubation with the primary antibody, sections were first incubated in the presence of a biotinylated secondary antibody (goat anti-rabbit, 1:200; Vector Laboratories, Cat. VEC-BA-1000) and then treated with avidin-biotin–horseradish peroxidase complex (ABC Elite kit, Vector Laboratories, Cat VEC-PK-6100). The brightfield signal was developed using a 3,3′-diaminobenzidine kit (Vector Laboratories Cat. VEC-SK-4100). Brighfield images were taken using an AxioScan.Z1-068 slide scanner, 20x/0.8NA objective. To assess microglia morphology from brain sections, images were processed as follows. First, background was subtracted using a rolling ball algorithm (radius of 25 pixels, light background) in Fiji (ImageJ version 2.1.0/1.53c), images were despeckled and Gaussian blur filter (sigma=0.5) was used. Iba1-positive cells were then identified from inverted, gray scale images in CellProfiler. Briefly, images were thresholded via a “Robust Background” method, with threshold smoothing factor 1.0 and smoothing scale 1.34. Different parameters describing microglia morphology (area of the cell body, perimeter, solidity, number of branches/cell, average branch length, triple point, and bounding box area) and the number of microglia were automatically determined. Data analysis was performed in R (version 4.2.0, R Core Team, Vienna, Austria).

### Real-time PCR (RT-qPCR)

RT-PCR analyses were performed as previously described ^21^. In brief, total RNA was extracted from microglial using a Direc-zol RNA Miniprep kit (Cat. R2050, Zymo Research), and cDNA synthesis was performed using Moloney murine leukemia virus reverse transcriptase (Cat. M3681, Promega) and random primers (Cat. C118A, Promega). A mass of 0.5 μg of RNA was denatured at 70 °C for 5 min in the presence of 0.75 μg of random primers in a 7.5 μL final volume. RT-PCR was carried out at 37 °C for 1 hour, and the enzyme was inactivated at 75 °C for 5 min. For each sample, control reactions were routinely performed without addition of reverse transcriptase. A 1:20 cDNA dilution was amplified using SYBR Green chemistry in triplicate in a 96-well plate using GoTaq qPCR Master Mix technology (Cat. A6001, Promega) according to the manufacturer’s protocol (5 μL of qPCR master mix, 0.15 μL of 100 mM primers each) and 4.7 μL of cDNA) using a StepOnePlus Real-Time PCR (Thermo Fisher Scientific) with the following thermal profile: 2 min at 95 °C and 40 cycles of 15 s at 95 °C and 1 min at 60 °C. The primers used are listed in Table 1 (Eurofins), and quantification was performed using the comparative CT method (2−ΔΔCt).

**Table 1:**
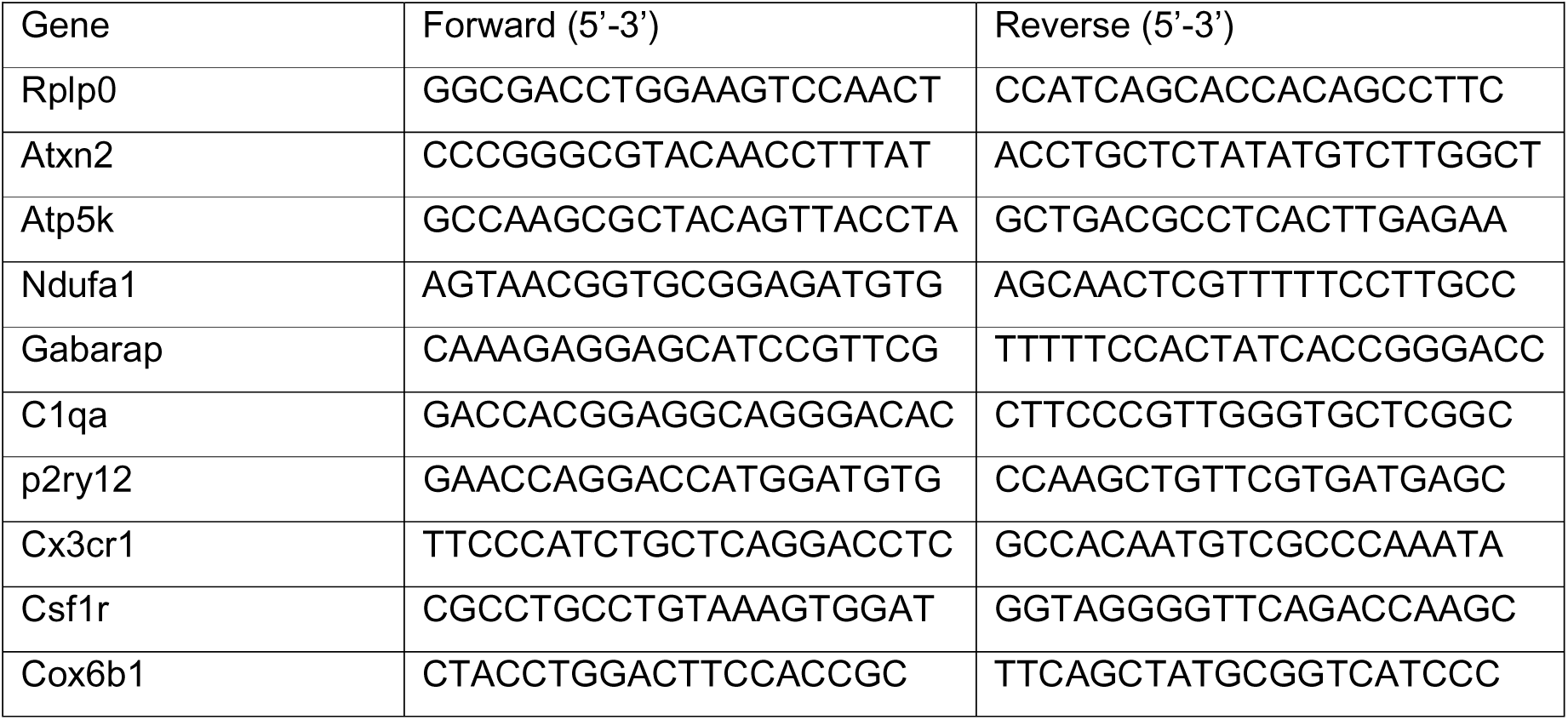
Primers used for RT-PCR.

### Metabolomic analysis

Metabolite extraction was performed by adding a mixture of MeOH/ACN/HLJO (5:3:2) containing an internal standard (IS) mix (1 µg of (Tryptophan(indole)-d5; Phenylalanine-15N; Stearic Acid-d3) to each sample, ensuring a final volume of 300 µL of extraction buffer per 3 x 10LJ cells as part of the normalization step. The mixture was vortexed for 10 seconds after breaking the cellular membrane by 2 freeze and thaw cycles (2 min for each cycle) using ethanol and dry ice cold bath and subjected to ultrasonic agitation for 30 minutes at 4°C. Afterward, the samples were centrifuged at 14,000 g for 20 minutes at 4°C. Supernatants were transferred to another tube and dried under a nitrogen atmosphere. Finally, the dried residues were resuspended in 120 µl H_2_O/ACN 1:1 and centrifuged again at 14,000 g for 15 minutes at 4 °C. 15 µL of the supernatant were injected into the LC-MS/MS system for analysis. Quality control (QC) samples were prepared as pool of all the samples, by mixing equal volume of each extracted sample and analyzed within the queue. All samples were analyzed in technical duplicate.

The metabolomics profiling was performed on UPLC (UPLC 1290 system, Agilent Technologies) coupled to mass spectrometer (TripleTOF 5600+, SCIEX) equipped with an electrospray ionization source (ESI), operating in both positive and negative ionization modes. Chromatographic separation was achieved on a SeQuant® ZIC®-pHILIC 5µm, 150 x 2.1 mm polymer capillary column. A gradient of solvent A (acetonitrile) and solvent B (water containing 20 mM (NHLJ)LJCOLJ and 0.1% ammonia) was used for separation at a flow rate of 200 µL/min. The gradient started with 20% solvent B for 2 minutes, then increased linearly to 80% solvent B over the next 15 minutes. It was rapidly adjusted back to 20% of solvent B within 10 seconds and held at this composition for equilibration until the end of the run (22 minutes). The column temperature was maintained at 45°C, while the autosampler was kept at 4°C to ensure optimal performance and sample stability. Full scan spectra were acquired in the mass range from m/z 75 to 1000 with a SWATH modality acquisition of 20 windows. The source parameters were: Gas 1: 33 psi, Gas 2: 58 psi, Curtain gas: 35 psi, Temperature: 500 °C and ISVF (IonSpray Voltage Floating): 5500 V (-4500 V for negative polarity), DP: 80 V, CE: 35 V with a spread of 15V. Data processing and targeted metabolite identification were performed using SCIEX OS software. Metabolites were identified based on high-resolution parent ion detection and corresponding mass fragmentation spectra. For confident identification, the metabolites of interest were annotated by comparing MS spectra and retention times with those of reference standards. The analyzed compounds were associated with key energy metabolism pathways, including glycolysis (glucose, glucose-6-phosphate, fructose-6-phosphate, glyceraldehyde-3-phosphate, 3-phosphoglycerate, phosphoenolpyruvate, and lactate), the pentose phosphate pathway (ribose-5-phosphate and 6-phosphogluconate), the tricarboxylic acid (TCA) cycle (citrate, isocitrate, succinate, fumarate, and malate), and glutaminolysis (glutamine, glutamate, and α-ketoglutarate) and glutathione.

### Library preparation, RNA sequencing and data analysis

Total RNA was extracted from microglia using the Direct-zol RNA Miniprep Kit (Cat. R2051, Zymo Research), following the manufacturer’s protocol. A total of eight mice were used per treatment group. RNA from four mice was pooled to generate two pools per treatment group. RNA quantity and quality were assessed using the Tapestation 4200 System (Agilent, Santa Clara, CA, USA) and the NanoDrop spectrophotometer (Thermo Fisher Scientific, Pittsburgh, PA, USA). RNA samples with RIN >7 were selected for downstream processing. Libraries were prepared using the TruSeq Stranded mRNA Prep Kit (Illumina, San Diego, CA, USA) following the manufacturer’s protocol. Starting with 500 ng of total RNA, mRNA was purified using poly-A selection and fragmented. Reverse transcription was performed to generate first-strand cDNA, followed by second-strand synthesis to produce double-stranded cDNA. The cDNA molecules were then subjected to 3’-end adenylation, adapter ligation, and cleanup steps for further sequencing. Libraries were amplified using 12 PCR cycles. Paired-end sequencing was carried out on the NextSeq 550 instrument (Illumina) achieving an average output of 50 million high-quality reads per sample.

Raw RNA-seq reads in FASTQ format were pre-processed for quality control using FastQC (v0.11.9). Adapter sequences and low-quality bases were removed using Trim Galore (v0.6.7) with default parameters. After trimming, reads were aligned to the Mus musculus reference genome (mm10, GRCm38) using HISAT2 (v2.2.1) with the following parameters: --dta --rna-strandness RF --no-mixed --no-discordant -p 8. Generated SAM files were converted into BAM format with SAMtools (v1.15.1) and then processed with RSEM (v1.3.3) to quantify gene expression. Gene-level raw counts were obtained using RSEM’s rsem-calculate-expression with the --paired-end and --bam options enabled. For downstream normalizations, the gene-level counts were imported to R and subsequently processed by the package EdgeR (Bioconductor version 3.14). Gene-length and sequencing-depth-normalized values of RPKM were obtained by using the rpkm function from the EdgeR. The quantification of TPM values was performed using Salmon^58^ . Differential expression analysis was performed using limma-voom^59^ on the Galaxy platform (https://usegalaxy.eu/?tool_id=toolshed.g2.bx.psu.edu%2Frepos%2Fiuc%2Flimma_voom%2Flimma_voom%2F3.58.1%2Bgalaxy0&version=latest; task started 2023-01-17). Limma (Linear Models for Microarray Data) was employed ^60^ and the voom method^61^ was used to apply precision weights for RNA-seq read counts. Differential expression analysis was carried out by comparing the vehicle vs. CBE datasets. No weights were applied to the samples, and eBayes with robust settings were utilized for the analysis. Subsequently, differential expression of upregulated and downregulated genes with a p-value < 0.05 was analyzed using Cytoscape and the ClueGO plugin (ClueGO v2.5.9). The Enrichment/Depletion (Two-sided hypergeometric test) was used for statistical analysis, and Bonferroni step-down correction was applied. The ontology used were KEGG_25.05.2022, GO_CellularComponent-EBI-UniProt-GOA-ACAP-ARAP_25.05.2022_00h00, and GO_BiologicalProcess-EBI-UniProt-GOA-ACAP-ARAP_25.05.2022_00h00. Minimum GO levels of 3 and a maximum GO level of 8 were set with a kappa score threshold of 0.4.

### Statistical analysis

Unless otherwise indicated in the figure legend, variables are presented as the mean with standard deviation. Statistical analyses were performed using Prism 7 (Version 7.00, GraphPad Software Inc.). T-tests were used to determine if there were significant differences in means between two groups. One-way ANOVA was used to determine if there were statistically significant differences in means among three or more independent groups; post hoc Tukey’s test was used to compare every mean with every other mean, or Dunnett’s test was used to compare every mean to a control mean. Two-way ANOVA followed by Sidak’s post hoc test was used to determine if the responses were affected by two factors in a multiple comparison. A p-value lower than 0.05 was considered to indicate statistical significance.

## Acknowledgements

The authors are grateful to the financial support from the EU Joint Programme - Neurodegenerative Disease Research (JPND) project (GBA-PaCTS, 01ED2005B and GBA-PARK n. 212), PNRR M4C2-Investimento 1.4-CN00000041-PNRR_CN3RNA_SPOKE9 (to P.C.) and Dipartimento DISS, Linea 2, Università degli Studi di Milano (to E.B.)

## Author contributions

Conceptualization: P.C., E.B., and A.V.; methodology: E.B., E.M.S, L.V.P., D.D., P.R., L.F.; Investigation: E.B., A.V., A.P., D.T., S.V., L.V.P., D.D., P.R., L.F, O.R., M.T., E.M.S., C.M. ; formal analysis: E.B., P.R,; Funding acquisition: P.C., D.A.D.M.; supervision: P.C., D.D., M.M.,A.A.; writing of original draft: P.C., E.B., A.V., review and editing of the manuscript: P.C., D.D., A.A., L.F.; All authors read and approved the final manuscript.

## Additional information

**Correspondence** and requests for materials should be addressed to Paolo Ciana.

## Bibliography

1. Horowitz, M. et al. The human glucocerebrosidase gene and pseudogene: structure and evolution. Genomics 4, 87–96 (1989).

2. Beutler, E. Gaucher disease: new molecular approaches to diagnosis and treatment. Science. 256, 794–799 (1992).

3. Gan-Or, Z. et al. Differential effects of severe vs mild GBA mutations on Parkinson disease. Neurology 84, 880–887 (2015).

4. Migdalska-Richards, A. & Schapira, A. H. V. The relationship between glucocerebrosidase mutations and Parkinson disease. J. Neurochem. 139, 77–90 (2016).

5. Parnetti, L. et al. Cerebrospinal fluid lysosomal enzymes and alpha-synuclein in Parkinson’s disease. Mov. Disord. 29, 1019–1027 (2014).

6. Soria, F. N. et al. Glucocerebrosidase deficiency in dopaminergic neurons induces microglial activation without neurodegeneration. Hum. Mol. Genet. 26, 2603–2615 (2017).

7. Kettenmann, H., Uwe Karsten, H., Mami, N. & Alexei, V. Physiology of microglia. Physiol. Rev. 91, 461–553 (2011).

8. Sierra, A., Paolicelli, R. C. & Kettenmann, H. Cien Años de Microglía: Milestones in a Century of Microglial Research. Trends Neurosci. 42, 778–792 (2019).

9. Bernier, L.-P., York, E. M. & MacVicar, B. A. Immunometabolism in the Brain: How Metabolism Shapes Microglial Function. Trends Neurosci. (2020).

10. Bernier, L.-P. et al. Microglial metabolic flexibility supports immune surveillance of the brain parenchyma. Nat. Commun. 11, 1559 (2020).

11. Farfel-Becker, T. et al. Spatial and temporal correlation between neuron loss and neuroinflammation in a mouse model of neuronopathic Gaucher disease. Hum. Mol. Genet. 20, 1375–1386 (2011).

12. Vitner, E. B., Farfel-Becker, T., Eilam, R., Biton, I. & Futerman, A. H. Contribution of brain inflammation to neuronal cell death in neuronopathic forms of Gaucher’s disease. Brain 135, 1724–1735 (2012).

13. Rocha, E. M. et al. Sustained systemic glucocerebrosidase inhibition induces brain α-synuclein aggregation, microglia and complement C1q activation in mice. Antioxid. Redox Signal. 23, 550–564 (2015).

14. Vardi, A. et al. Delineating pathological pathways in a chemically induced mouse model of Gaucher disease. J. Pathol. 239, 496–509 (2016).

15. Boka, G. et al. Immunocytochemical analysis of tumor necrosis factor and its receptors in Parkinson’s disease. Neurosci. Lett. 172, 151–154 (1994).

16. McGeer, P. L., Itagaki, S., Boyes, B. E. & McGeer, E. G. Reactive microglia are positive for HLA-DR in the substantia nigra of Parkinson’s and Alzheimer’s disease brains. Neurology 38, 1285 (1988).

17. Richter, F. et al. A GCase chaperone improves motor function in a mouse model of synucleinopathy. Neurotherapeutics 11, 840–856 (2014).

18. Keatinge, M. et al. Glucocerebrosidase 1 deficient Danio rerio mirror key pathological aspects of human Gaucher disease and provide evidence of early microglial activation preceding alpha-synuclein-independent neuronal cell death. Hum. Mol. Genet. 24, 6640– 6652 (2015).

19. Mullin, S. et al. Brain microglial activation increased in glucocerebrosidase (GBA) mutation carriers without Parkinson’s disease. Mov. Disord. 36, 774–779 (2021).

20. Hayes, J. D. & Dinkova-Kostova, A. T. The Nrf2 regulatory network provides an interface between redox and intermediary metabolism. Trends Biochem. Sci. 39, 199–218 (2014).

21. Brunialti, E. et al. Inhibition of microglial β-glucocerebrosidase hampers the microglia-mediated antioxidant and protective response in neurons. J. Neuroinflammation 18, 1–18 (2021).

22. Rizzi, N. et al. In vivo imaging of early signs of dopaminergic neuronal death in an animal model of Parkinson’s disease. Neurobiol Dis. 114, 74–84 (2018).

23. Brunialti, E. et al. sex-specific microglial responses to glucocerebrosidase inhibition: relevance to GBA1-Linked Parkinson’s Disease. Cells 12, 343 (2023).

24. Zanier, E. R., Fumagalli, S., Perego, C., Pischiutta, F. & De Simoni, M.-G. Shape descriptors of the “never resting” microglia in three different acute brain injury models in mice. Intensive Care Med. Exp. 3, 0–18 (2015).

25. Liu, Y. et al. Mice with type 2 and 3 Gaucher disease point mutations generated by a single insertion mutagenesis procedure (SIMP). Proc. Natl. Acad. Sci. 95, 2503–2508 (1998).

26. La Vitola, P. et al. Mitochondrial oxidant stress promotes α-synuclein aggregation and spreading in mice with mutated glucocerebrosidase. *npj Park*. Dis. 10, 1–13 (2024).

27. Migdalska-Richards, A. et al. The L444P Gba1 mutation enhances alpha-synuclein induced loss of nigral dopaminergic neurons in mice. Brain 140, 2706–2721 (2017).

28. Bindea, G. et al. ClueGO: a Cytoscape plug-in to decipher functionally grouped gene ontology and pathway annotation networks. Bioinformatics 25, 1091–1093 (2009).

29. Hansen, G. E. & Gibson, G. E. The α-ketoglutarate dehydrogenase complex as a hub of plasticity in neurodegeneration and regeneration. Int. J. Mol. Sci. 23, 12403 (2022).

30. Liu, P.-S. et al. α-ketoglutarate orchestrates macrophage activation through metabolic and epigenetic reprogramming. Nat. Immunol. 18, 985–994 (2017).

31. Liu, S., Yang, J. & Wu, Z. The regulatory role of α-ketoglutarate metabolism in macrophages. Mediators Inflamm. 2021, 5577577 (2021).

32. Li, L. et al. Resolvin D1 reprograms energy metabolism to promote microglia to phagocytize neutrophils after ischemic stroke. Cell Rep. 2023; 42: 112617–112617. (2023).

33. Lundgaard, I. et al. Direct neuronal glucose uptake heralds activity-dependent increases in cerebral metabolism. Nat. Commun. 6, 6807 (2015).

34. Li, H. et al. Neurons require glucose uptake and glycolysis in vivo. Cell Rep. 42, (2023).

35. Ledderose, C. et al. The purinergic receptor P2Y11 choreographs the polarization, mitochondrial metabolism, and migration of T lymphocytes. Sci. Signal. 13, eaba3300 (2020).

36. Hu, Y. et al. mTOR-mediated metabolic reprogramming shapes distinct microglia functions in response to lipopolysaccharide and ATP. Glia 68, 1031–1045 (2020).

37. Nair, S. et al. Lipopolysaccharide-induced alteration of mitochondrial morphology induces a metabolic shift in microglia modulating the inflammatory response in vitro and in vivo. Glia 67, 1047–1061 (2019).

38. Cheng, J. et al. Early glycolytic reprogramming controls microglial inflammatory activation. J. Neuroinflammation 18, 129 (2021).

39. Gegg, M. E. et al. Glucocerebrosidase deficiency in substantia nigra of parkinson disease brains. Ann. Neurol. 72, 455–463 (2012).

40. Gegg, M. E., Menozzi, E. & Schapira, A. H. V. Glucocerebrosidase-associated Parkinson disease: Pathogenic mechanisms and potential drug treatments. Neurobiol Dis. 166, 105663 (2022).

41. Aflaki, E. et al. A characterization of Gaucher iPS-derived astrocytes: potential implications for Parkinson’s disease. Neurobiol Dis. 134, 104647 (2020).

42. Wang, L. et al. Neuronal activity induces glucosylceramide that is secreted via exosomes for lysosomal degradation in glia. Sci. Adv. 8, eabn3326 (2022).

43. Revel-Vilk, S., Szer, J. & Zimran, A. Hematological manifestations and complications of Gaucher disease. Expert Rev. Hematol. 14, 347–354 (2021).

44. Scheiblich, H. et al. Microglia rescue neurons from aggregate-induced neuronal dysfunction and death through tunneling nanotubes. Neuron 112, 3106–3125 (2024).

45. Cserép, C. et al. Microglia monitor and protect neuronal function through specialized somatic purinergic junctions. Science. 367, 528–537 (2020).

46. Navarro, E. et al. Dysregulation of mitochondrial and proteolysosomal genes in Parkinson’s disease myeloid cells. *Nat*. aging 1, 850–863 (2021).

47. Yang, S. et al. Microglia reprogram metabolic profiles for phenotype and function changes in central nervous system. Neurobiol Dis. 152, 105290 (2021).

48. Sanman, L. E. et al. Disruption of glycolytic flux is a signal for inflammasome signaling and pyroptotic cell death. Elife 5, e13663 (2016).

49. Aoyama, K. & Nakaki, T. Impaired glutathione synthesis in neurodegeneration. Int. J. Mol. Sci. 14, 21021–21044 (2013).

50. Osellame, L. D. et al. Mitochondria and quality control defects in a mouse model of Gaucher disease—links to Parkinson’s disease. Cell Metab. 17, 941–953 (2013).

51. Schapira, A. H. V & Tolosa, E. Molecular and clinical prodrome of Parkinson disease: implications for treatment. Nat. Rev. Neurol. 6, 309–317 (2010).

52. Bottani, E. et al. Therapeutic approaches to treat mitochondrial diseases:“one-size-fits-all” and “precision medicine” strategies. Pharmaceutics 12, 1083 (2020).

53. Xia, D., Liu, Y., Wu, P. & Wei, D. Current advances of mitochondrial dysfunction and cardiovascular disease and promising therapeutic strategies. Am. J. Pathol. 193, 1485–1500 (2023).

54. Bhullar, S. K. & Dhalla, N. S. Status of mitochondrial oxidative phosphorylation during the development of heart failure. Antioxidants 12, 1941 (2023).

55. Seo, D. R. et al. Cross talk between P2 purinergic receptors modulates extracellular ATP-mediated interleukin-10 production in rat microglial cells. Exp. Mol. Med. 40, 19–26 (2008).

56. Gendron, F. et al. P2X7 nucleotide receptor activation enhances IFNγ-induced type II nitric oxide synthase activity in BV-2 microglial cells. J. Neurochem. 87, 344–352 (2003).

57. Khakpay, R. et al. Potentiation of the glutamatergic synaptic input to rat locus coeruleus neurons by P2X7 receptors. Purinergic Signal. 6, 349–359 (2010).

58. Patro, R., Duggal, G., Love, M. I., Irizarry, R. A. & Kingsford, C. Salmon provides fast and bias-aware quantification of transcript expression. Nat. Methods 14, 417–419 (2017).

59. Smyth, G. K. Limma: linear models for microarray data. in Bioinformatics and computational biology solutions using R and Bioconductor 397–420 (Springer, 2005).

60. Gentleman, R., Carey, V., Huber, W., Irizarry, R. & Dudoit, S. Gentleman, et al., 2005. (Springer Science & Business Media, 2005).

61. Law, C. W., Chen, Y., Shi, W. & Smyth, G. K. voom: Precision weights unlock linear model analysis tools for RNA-seq read counts. Genome Biol. 15, 1–17 (2014).

