## supplementary for "Metabolic reprogramming and altered ATP content impair neuroprotective functions of microglia in β-glucocerebrosidase deficiency models"

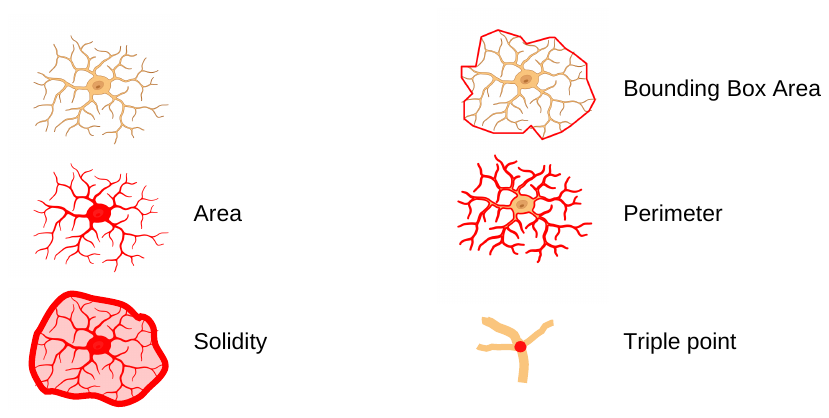

**Figure Supplementary 1:** the red color indicates the quantified morphological value reported in Figure 1. Solidity represents the ratio between the object's area and its convex area, while the number of triple points refers to junctions where exactly three branches meet.

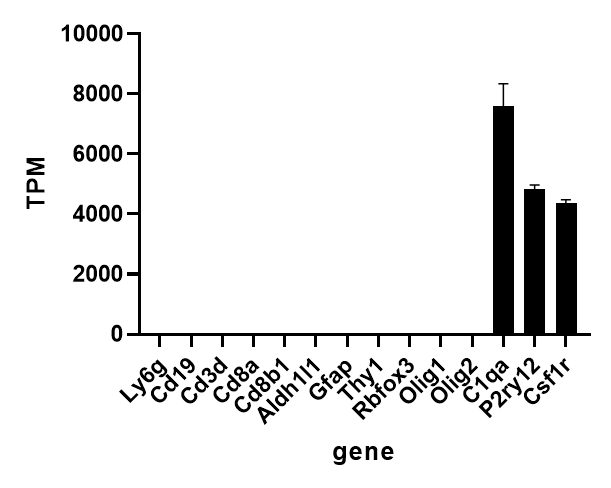

**Figure Supplementary 2:** Analysis of the TPM values obtained from the transcriptomic assay described in Figure 2 highlights that the samples were highly enriched in microglia, as microglial genes (*C1qa, P2ry12, Csf1r*) are drastically overrepresented compared to genes of other brain cells.

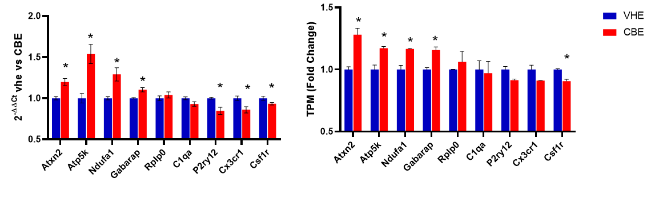

**Figure Supplementary 3:** qPCR validation(left): A total of 9 genes were selected. Data are expressed using the 2^-ΔΔCt^ method. Columns represent the mean ± SEM of 8 animals per experimental group. *p < 0.05; determined by unpaired t-test comparisons. The expression analyses of the selected genes gave results consistent with those obtained by RNAseq (right).

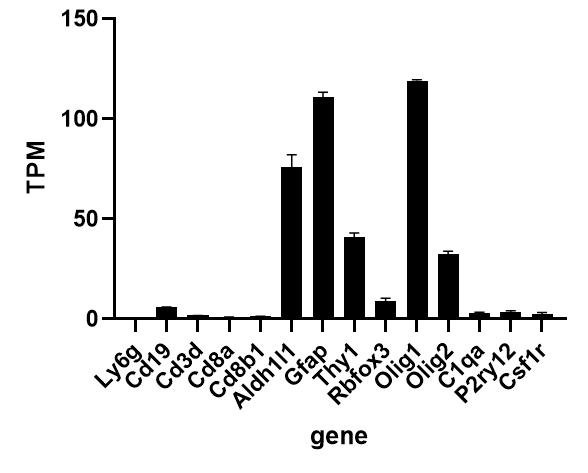

**Figure Supplementary 4:** Gene expression analysis of the cellular brain population from which the microglia were removed highlights that the sample is enriched in neuron, astrocyte, and oligodendrocyte transcripts.

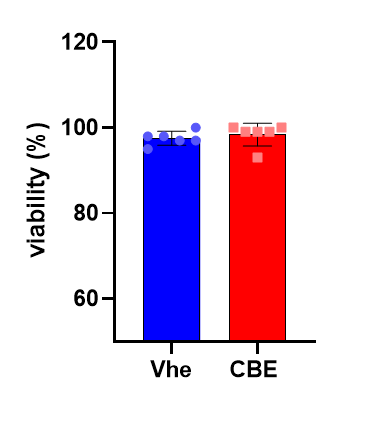

**Figure Supplementary 5:** Viability assay using trypan blue in cocultures of SK-N-BE and BV-2 cells treated with vehicle or 200 µM CBE for 48 hours. Data represent percentage viability relative to vehicle-treated cells ± SEM (n=6); no significant difference identified with unpaired t-test.

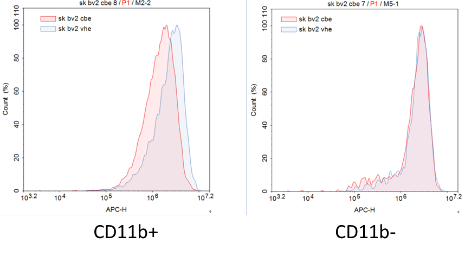

**Figure Supplementary 6:** Representative graph of the FACS analysis reported in Figure 4D showing mitochondrial staining intensity in CD11b+ and CD11b- populations following treatment with 200 µM CBE for 48 hours.

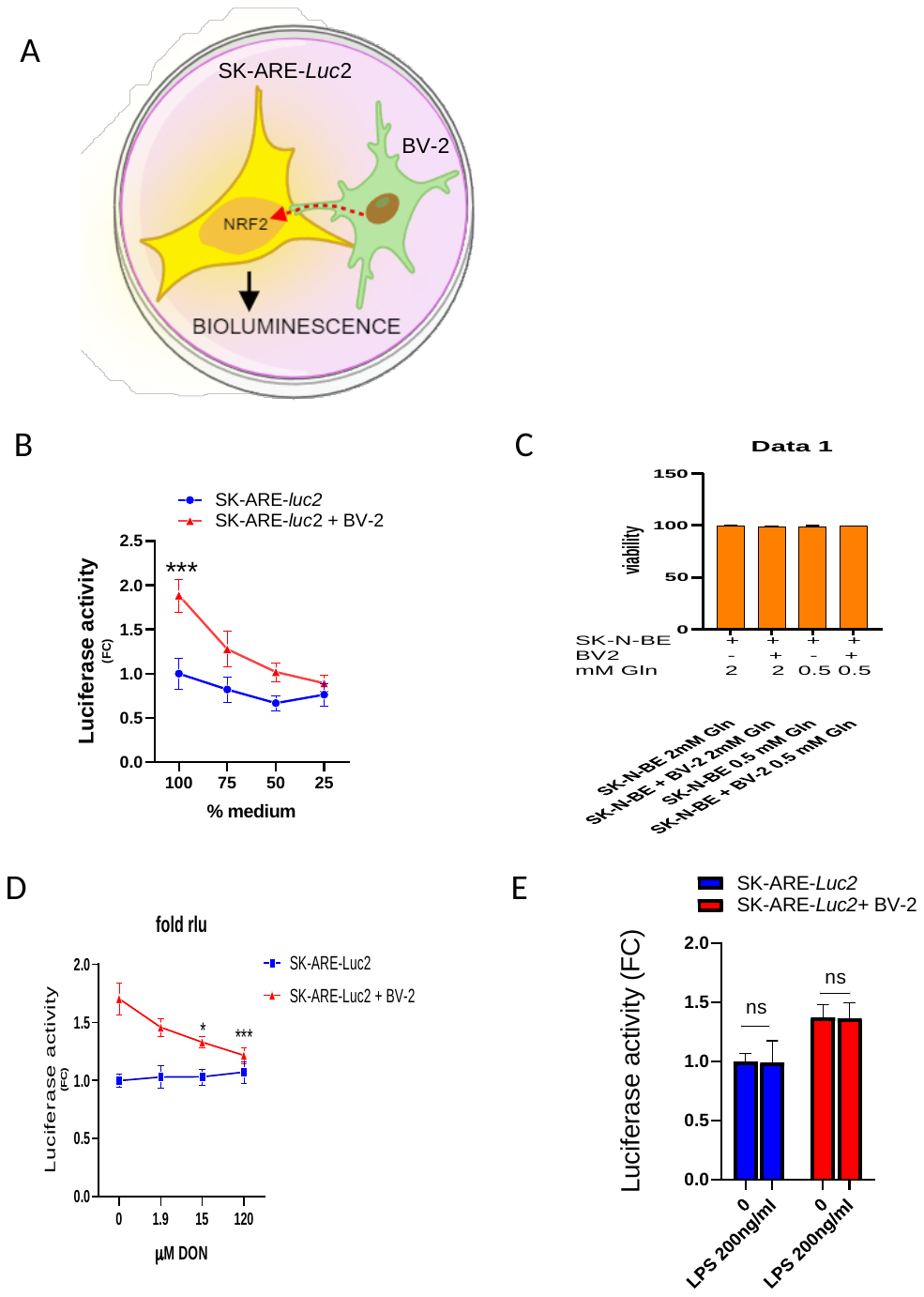

**Supplementary 7:**

(A) Schematic representation of co-culture experiments designed to evaluate the effects of various treatments on microglia-neuron communication. The microglia are capable of activating the NFE2L2 (NRF2) transcription factor in neurons, a mechanism that can be detected using the SK-N-BE bioluminescent cell reporter for NFE2L2 (SK-ARE-*luc2*).

activation in SK-ARE-*luc2*/BV-2 cocultures.

(B) Luciferase activity measured in protein extracts derived from SK-ARE-*luc2* cells or coculture of SK-ARE-*luc2* and BV-2 cells grown for 6 hours in complete media or media diluted in HBSS. Data represents fold change (FC) of luciferase activity versus SK-ARE-*luc2* grown in complete media ± SEM (n=6); ***p<0.001 versus SK-ARE-*luc2* calculated with one-way ANOVA followed by Sidak’s multiple comparisons test.

(C) Viability assay on cells grown with reduced amounts of glutamine highlights that there is no viability deficit. SK-N-BE and BV-2 cells were grown for 6 hours with different concentrations of glutamine.

(D) Inhibition of multiple enzymes that use glutamine through the mimetic 6-diazo-5-oxo-L-norleucine (DON) reduces microglia-to-neuron communication. SK-ARE-*luc2* cells or coculture of SK-ARE-*luc2* + BV-2 were treated with different doses of DON for 6 hours, and the luciferase activity was measured in cellular extracts. Data represents fold change (FC) of luciferase activity versus SK-ARE-*luc2* ± SEM. *p<0.05, **p<0.01, calculated with one-way ANOVA followed by Sidak’s multiple comparisons test versus vehicle-treated cells.

(E) Luciferase activity measured in protein extracts derived from SK-ARE-*luc2* cells or coculture of SK-ARE-*luc2* and BV-2 cells treated for 4 hours with 200 ng/ml of LPS. Data represent fold change (FC) of luciferase activity versus SK-ARE-*luc2* grown in complete media ± SEM (n=7); ns, not significant, calculated with one-way ANOVA followed by Sidak’s multiple comparisons test.
